# Translating electrophysiological signatures of awareness into thalamocortical mechanisms by inverting systems-level computational models across arousal states

**DOI:** 10.1101/2023.10.11.561970

**Authors:** Vicente Medel, Eli J. Muller, Brandon R. Munn, Cameron Casey, Robert D. Sanders, James M. Shine

## Abstract

While consciousness never fades during wakefulness, there is a paradoxical coexistence of consciousness during dreaming states. It’s also possible for sensory awareness to be either present or absent when awakened from seemingly-identical states of sedation and anaesthesia. Traditionally, these states have been characterised in terms of their electroencephalographic neural correlates, however, without clear underlying neurobiological mechanisms. To bridge this gap, we invert a validated neural mass model of the corticothalamic system using scalp EEG collected during nonlinear transitions in conscious experience and sensory awareness across varying depths of dexmedetomidine sedation. We found that a decline in conscious experience and sensory awareness with dexmedetomidine was associated with a decrease in the engagement of excitatory thalamocortical loop resonances, along with an increase in inhibitory intrathalamic loop gains. These findings shed light into the neural mechanisms of conscious experience and sensory awareness, and explain how it fades across arousal states, bridging the gap between the neural correlates of consciousness and its underlying systems-level thalamocortical mechanisms.

## Introduction

Every waking moment is accompanied by a seemingly endless flow of experiences, with consciousness receding during certain stages of sleep and anaesthesia. Indeed, subjects roused from sleep or light anaesthesia may in some occasions report detailed and vivid internal dreaming experiences or even external sensory awareness, whereas in other seemingly similar states, they are utterly devoid of experience when roused (Stickgold et al., 2001; Nir & Tononi, 2010; Siclari et al., 2017; Kotsovolis & Komninos, 2009; Noreika et al. 2011; Sanders et al., 2012; Sanders et al., 2017). How these different states arise in the brain is an open question. From waking states, we know that subjects dynamically shift their brain activity through a variety of states that regulate internally- and externally-driven awareness of the world (McGinley et al., 2013; McCormick et al., 2020; Munn et al., 2021). These fluctuations are thought to arise through distinct activity patterns in a distributed set of thalamocortical circuits that modulate the brain’s intrinsic activity and its responsiveness to external stimuli (Kosciessa et al., 2021; Shine et al., 2023). Although there is evidence of the role of thalamocortical circuits in shaping brain dynamic underlying changes from awake to unconscious sleep or anaesthesia (Redinbaugh et al., 2020; Tasserie et al., 2022; Muller et al., 2023), it is still unknown how this system relates to the rather paradoxical occurrence of both presence and absence of experience within the same behavioural state.

Decades of studies have revealed tractable EEG neural markers that effectively differentiate various states of arousal, each tied to potential (but unconstrained) neurobiological mechanisms (Lendner et al., 2020; Gao et al., 2017; Trakoshis et al., 2020). Notably, neurophysiological measures like alpha power or 1/f spectral exponent activity can distinguish between arousal states and their transitions (Podvalny et al., 2021; Pfeffer et al., 2022; Medel et al., 2021; Gao et al., 2017; Stitt et al., 2018; Yüzgec et al., 2018; Nestvogel & McCormick, 2022; Lendner et al., 2020). While these methodologies provide invaluable perspectives on the interplay between brain electrophysiological activity and conscious awareness, there remains a notable gap in how these processes align with the intricate neurobiological architecture of the brain underlying consciousness and its emergence.

To navigate this impasse, we invert a well-validated neural mass model of the corticothalamic system, grounded in physiological and physical principles, using the BrainTrack method (Abeysuriya et al., 2015; Abeysuriya et al., 2016) to translate spectral EEG experimental measurements into model parameters characterising neural populations and their thalamocortical connection gains. We blend this method with a dataset exploring non-linear transitions of the presence and absence of conscious experience and sensory awareness undering sedation (Casey et al., 2022). By using a novel corticothalamic model-driven data analysis strategy, we here interrogate the underlying neurobiological mechanisms of conscious experience and sensory awareness across arousal states.

## Results

We collected 256-channel scalp EEG from 20 participants enrolled in the Understanding Consciousness Connectedness and Intra-Operative Unresponsiveness Study (NCT03284307) to undergo the Serial Awakening Paradigm (Casey et al., 2022). Each participant received a titrated dose of dexmedetomidine, which is an alpha-2 adrenergic receptor agonist that increases auto-inhibition of the locus coeruleus, thus depriving the brain of the noradrenergic tone required to support consciousness (Carter et al., 2010; Berridge et al., 2012) and trigger arousal (Sara & Bouret, 2012). 2-10 minute samples of whole-scalp EEG were recorded prior to wakening, and states were classified based on the subject’s experience described after a brief structured interview consisting of questions designed to assess if the participant had been having a conscious experience directly before the name call and if the experience was connected to the environment through the senses. Four groups were identified: Wake (N=20); connected consciousness (CC; N = 16); disconnected consciousness (DC; N = 20) and no experience (NE; N = 20). Although subjects were always rousable (a known feature of dexmedetomidine sedation (Casey et al., 2022; Sanders & Maze, 2011), their conscious experiences were different in the three non-awake categories: in CC, subjects had been given the drug, were sedated but were aware of their environment; in DC, subjects reported dream reports on rousing; and in NE, subjects reported no awareness of the external world or dreams, upon rousing. We were able to use these distinct states as lenses through which to further interrogate our data.

To this end, we first analysed measures of brain state from scalp EEG to characterise the functional transitions across arousal states. After spectral parametrization of the signal (Donoghue et al., 2020), we obtained and characterised two distinct measures: the alpha relative power (alpha RP) and the 1/f exponent, each of which has been previously linked to changes in brain state (Podvalny et al., 2021; Pfeffer et al., 2022; Medel et al., 2021; Gao et al., 2017; Stitt et al., 2018; Yüzgec et al., 2018; Nestvogel & McCormick, 2022; Lendner et al., 2020). We found that both measures changed significantly across states (Fig. 1). Importantly, the EEG brain signal in Wake had a spatially distributed lower (i.e., flatter) 1/f exponent, while alpha RP were higher at occipital electrodes (1/f exponent Fz Wilcoxon p=0.0006, 1/f exponent Oz Wilcoxon p=0.041; alpha RP Fz Wilcoxon p=0.6, alpha RP Oz Wilcoxon p=0.003), which suggested that 1/f exponent had stronger effect in frontal while alpha RP only on occipital areas (Fig 1B, E, F; Supplementary Figure 1A). Although the topography of the 1/f slope and alpha RP were spatially correlated (Wake Spearman rho=0.92, p-value=4.01e-63; CC Spearman rho=0.47, p-value=7.29e-10; DC Spearman rho=0.13, p-value=0.11; NE Spearman rho=0.4, p-value=1.08e-08; Fig 1D; Supplementary Figure 1B), they have been proposed as separated biomarkers of brain state (Medel et al., 2021; Gao et al., 2017; Trakoshis et al., 2020), with separated hypothesised mechanisms and different neurophysiological correlates, and are calculated separately from the periodic and aperiodic components of the EEG signal (Donoghue et al., 2020). These results together with the prior evidence suggest both signal components representing brain states might be regulated by a systems-level mechanism.

**Figure 1.**
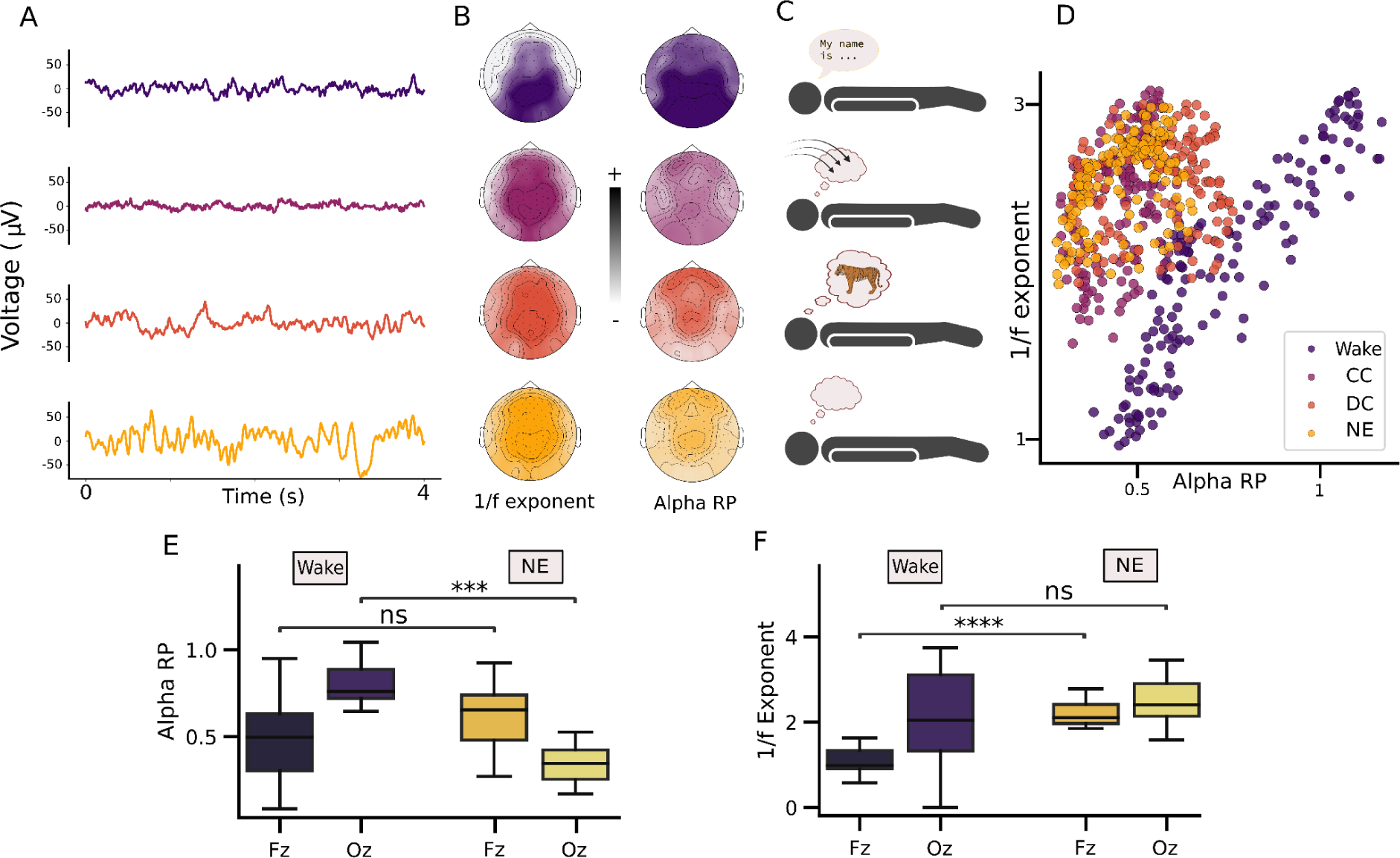
EEG activity across arousal states. (**A**) EEG signal traces for Wake (dark purple), Connected Consciousness (CC, light purple), Disconnected Consciousness (DC, orange), and No Experience (NE, yellow). (**B**) Left, shows the topography of the 1/f slope. Right, shows the topography of the Alpha Relative Power (RP). Both topographic plots have their colormaps in terms of brightness. (**C**) Denotes each arousal state. (**D**) Scatterplot with the same colour scheme representing the heterogeneity of the relation between 1/f slope and alpha RP across arousal states. (**E**) depicts the difference in alpha RP under Wake and NE, for Fz and Oz electrodes. (**F**) same as **E** but for the 1/f Exponent.

Based on previous work (Muller et al., 2023; Shine et al., 2023; Washke et al., 2021; Kocsiessa et al., 2021), we hypothesized that the changes in brain state were related to altered interactions between the thalamus and cerebral cortex, however the surface-based nature of the EEG signal makes it inherently difficult to test the predictions of this hypothesis. To circumvent this issue, we leveraged an established neural field corticothalamic model (Rowe et al., 2004, Muller et al., 2020; Rennie et al., 2002; Robinson et al., 2001, 2003; Abeysuriya et al., 2015, 2016) using only systems-level corticothalamic parameters, this model is able to reproduce a wide range of brain state characteristics, such as alpha rhythm (Robinson et al., 2003, O’Connor and Robinson, 2004) and 1/f slope (Abeysuriya & Robinson 2016), among other spectral properties seen in experimental EEG. To fit the data in the model, we calculated the power spectral density (PSD) of the Fz and Oz electrodes (see Methods for details) and then used the BrainTrack algorithm (Abeysuriya et al., 2015, 2016) to invert these data to the most plausible parameters in the neural field model of the corticothalamic system at each electrode across arousal states (see Methods, Figure 2). This allowed us to translate the brain state signatures identified from scalp EEG into the thalamocortical mechanisms that could support their expression, which in turn allowed us to directly test our hypothesized mechanism of altered consciousness as a function of dexmedetomidine sedation.

**Figure 2.**
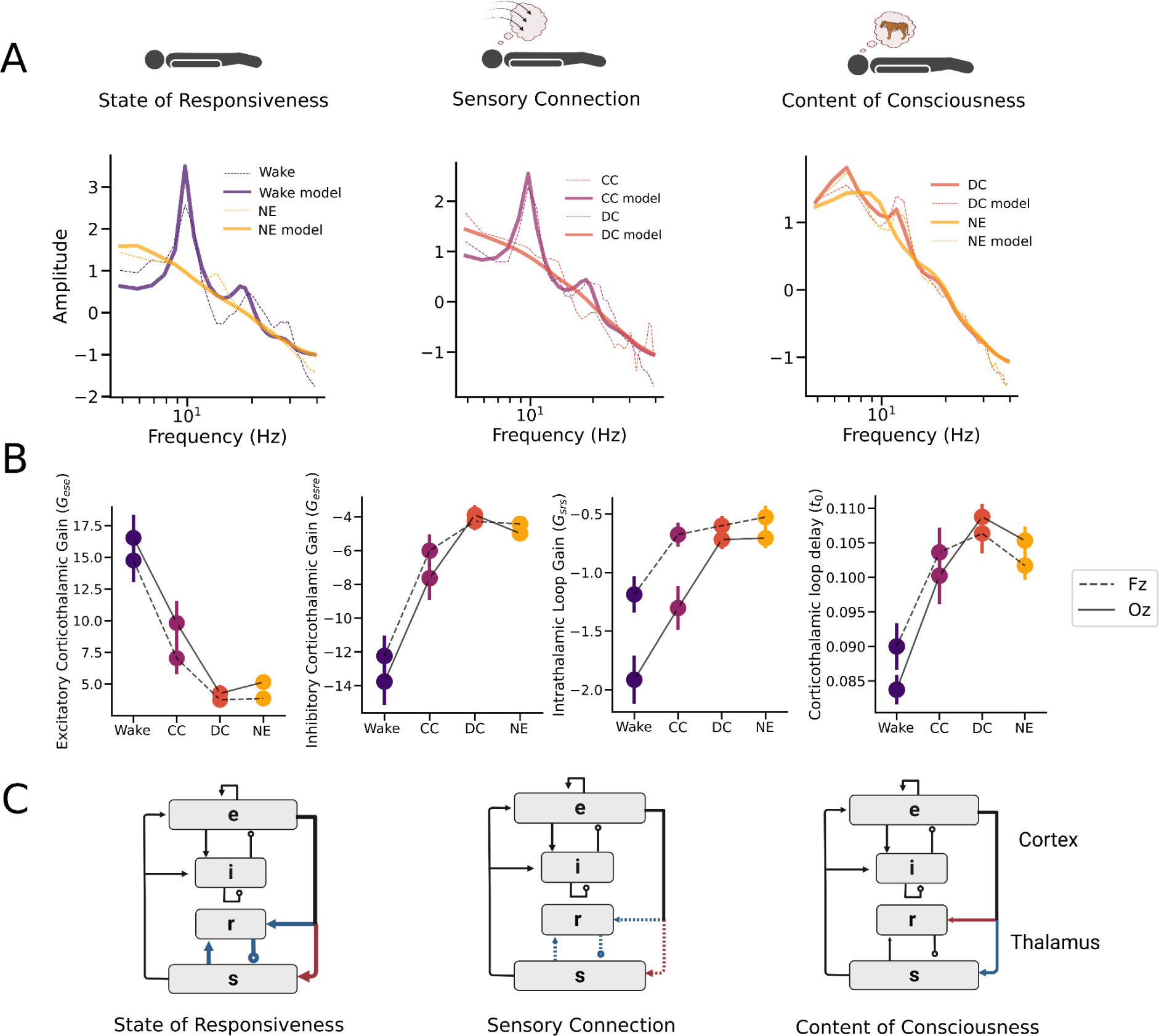
(**A**) Example of experimental EEG (dashed line) and the corresponding model fit (solid line) for each isolated arousal state comparison. (**B**) (**C**) schematic diagram of the modulated corticothalamic neural field model for each state comparison shown in **A**. The neural populations shown are cortical excitatory *e* and inhibitory *i*, thalamic reticular *r* and thalamic relay nuclei *s*. Red lines represent decreased gain modulation, while blue lines represent increased gain modulations.

The responsiveness comparison (Fig. 2A, Wake vs. NE) showed striking changes in shared parameters for both Fz and Oz for global corticothalamic parameters such as a decrease in excitatory (Fig 2A, bottom; G_ese_ ; Wilcoxon, Fz p-value=0.00017, Oz p-value=0.00012; Supplementary Figure 2) and increase in inhibitory (Fig 2A, bottom; G_esre_ ; Wilcoxon, Fz p-value = 0.0001, Oz p-value = 3.43e-05, Supplementary Figure 2) corticothalamic gains, as well as increase in intrathalamic gain (G_srs_ ; Wilcoxon, Fz p-value= 0.006, Oz p-value= 3.43e-05; Supplementary Figure 2) and the corticothalamic loop delay, t0 (Wilcoxon, Fz p-value=0.0063, Oz p-value=3.41e-05). To compare if these neural field model parameters are related to the changes of alpha RP and 1/f exponent, we correlated the significant parameters to the experimental spectral measures (Fig 1). We found a significant correlation between frontal 1/f exponent and G_ese_ (rho=-0.43, p=0.007), G_esre_ (rho=0.48, p=0.006), and t0 (rho=0.39, p=0.02), but not for occipital electrodes. In contrast, alpha RP was correlated with frontal t0 (rho=-0.37, p=0.03) and occipital G_srs_ (rho=-0.41, p=0.017). This suggests that the balance between the excitatory thalamic projections and the inhibitory reticular nucleus is a key determinant of the state of responsiveness differences, with decreasing corticothalamic loop delay (Fig. 2C), both mechanisms putatively underlying 1/f exponent and alpha RP consciousness biomarkers.

The impact of anaesthetic agents is actually highly nuanced, such that individuals, when woken, can sometimes report full experience of the world (which we label ‘connected consciousness’; or CC; Casey et al., 2022), whereas other times they either report dream-related experiences (‘disconnected consciousness’; or DC), or nothing at all (i.e., no experience; NE). Importantly, we were able to use our model inversion approach to map the differences between these states - which themselves characterise distinct signatures of consciousness - directly to thalamocortical loop gains (Figure 2A). Specifically, we defined the State of Responsiveness as Wake vs NE; Sensory Disconnection as comparing connected consciousness (CC) versus disconnected consciousness (DC); and the Content of Consciousness as the difference between CC vs NE, to compare experience in absence of sensory connection with unconscious state.

Next, we compared internally- and externally-induced low arousal states defined by Sensory Disconnection (Fig. 2B; CC vs. DC). These states are notoriously challenging to discriminate (Casey et al., 2022). Interestingly, we found changes consistent with those observed in the State of Responsiveness, albeit with a lower magnitude. Importantly, the changes were exclusively in occipital and not frontal electrodes. We found that sensory disconnection was related to a significant decrease in excitatory (G_ese_ ; Wilcoxon, Fz p-value=0.082, Oz p-value=0.0064), and an increase in inhibitory (G_esre_ ; Wilcoxon, Fz p-value=0.238, Oz p-value=0.0045) corticothalamic gain, increase in intrathalamic loop gain (G_srs_ ; Wilcoxon, Fz p-value= 0.362, Oz p-value= 0.0043), but no significative difference in corticothalamic loop delay, t0.

Studying the brain mechanisms underlying consciousness *per se* (i.e. the presence of phenomenological content) isolated from the perception of the external world has remained challenging, and there is no consensus whether it follows similar neural mechanisms as Sensory Disconnection or State of Responsiveness. To interpret the Content of Consciousness while controlling by Sensory Disconnection, we contrasted arousal states that reported sensory disconnection while being conscious against unconsciousness (DC vs NE). Surprisingly, we found that occipital electrodes showed a significant increase in excitatory (G_ese_ ; Wilcoxon, Fz p-value=0.00017, Oz p-value=0.0123) and decrease in inhibitory (Fig 2B, bottom; G_esre_ ; Wilcoxon, Fz p-value = 0.0001, Oz p-value = 0.0119) corticothalamic gains, as opposed to the tendency shown in the previous comparisons (Fig 2B), which suggest a non-linear change. Thus, we find that corticothalamic gain shapes arousal states with and without consciousness- and sensory-awareness, with a continuous yet non-monotonic decoupling between the cortex and the thalamus that coincided with an increasing intrathalamic loop gain. These results provide supporting evidence for our hypothesis that abnormal thalamocortical interactions are responsible for distinct brain states induced by anaesthesia.

## Discussion

Here, we first approach a spectral characterization of scalp EEG during the novel Serial Awakening Paradigm (Casey et al., 2022), and found that 1/f slope and the relative power of alpha oscillation were both related to the State of Responsiveness (Fig. 1E). Interestingly, both measures strongly covaried across the scalp (Fig 1D), but had spatially distinct predictive power on the state of the subjects (Fig 1B). Moreover, these spectral measures were not distinct in more fine-grained phenomenological differences across arousal states, such as Sensory Disconnection (CC vs DC) nor Content of Consciousness (DC vs NE). Due to different mechanisms putatively involved in 1/f slope (Gao et al., 2017, Trakoshis et al., 2020, Medel et al., 2021) and the relative power of alpha oscillations (Adam et al., 2023; Vijayan et al., 2013; Soplata et al., 2017), we leveraged an existing corticothalamic neural field model (Abeysuriya et al., 2015, 2016) to differentiate the common neural mechanisms underlying the distinct modes of consciousness in the human brain that change as a function of arousal. Across these states, we found that changes in arousal were associated with diminished thalamocortical resonance, which dropped due to a switch in the balance between excitatory and inhibitory processing within the thalamus. We also found that the State of Responsiveness was best explained by a decrease of excitatory corticothalamic gain, with an increase in inhibitory corticothalamic and intrathalamic gain, while these two structures being continuously decoupled, as measured by thalamocortical delay (t0). These mechanisms followed the same trend for Sensory Disconnection (CC vs DC), which suggests shared underlying mechanisms. Interestingly, we show an opposite change for Content of Consciousness (DC vs NE) consistent with non-linear biological changes prior to absence of experience (Pal et al., 2019; Boncompte et al., 2021; Cardone et al., 2023). Finally, we show that these same measures covaried with standard measures of EEG processing, such as the 1/f slope and power in the alpha band. Together, our approach helps to further our understanding of how neural activity and connectivity patterns change as a function of arousal.

Our results speak to the debate regarding whether consciousness is best mapped onto neural activity in the “front” or“back” of the cerebral cortex (Boly et al., 2017). Although Wake vs NE showed a frontal and occipital change in global thalamocortical gain, our analysis revealed that posterior and not frontal electrodes had a significant decoupling between the cortex and the thalamus that was informative of experience itself (Fig 2), discriminating both sensory-driven and internally-driven experiences, in close consistency with previous conscious correlates previously reported (Siclari et al., 2017, Casey et al., 2022). In addition, our approach further demonstrates the importance of interpreting indirect signatures of a complex, multi-scale system as selective evidence for effects at the site of measurement. In our case, even though surface EEG signatures were distinct across brain states, our modelling approach clearly shows that these signatures are attributable to mechanisms deep to the cortex - namely, in the highly-interconnected subcortical thalamic hub that is known to coordinate and shape whole brain dynamics (Bell & Shine, 2016; Shine et al., 2019; Muller et al., 2020; Shine et al., 2021; Muller et al., 2023). Our results thus add to the ongoing debate, while also expanding its scope to include descriptions of the brain that incorporate important regions outside the cerebral cortex in the explanation of changes in whole brain function.

Our results also enrich an evolving literature that highlights the key role of the thalamus and its connections with the cerebral cortex in supporting the neural states associated with waking consciousness. The thalamus has long been implicated in controlling conscious states (Steriade et al., 1993; ; Llinas et al., 1998; Seth et al., 2005; Shine et al., 2023), and recent empirical studies have helped to underscore these theoretical claims with robust neuroimaging data (Redinbaugh et al., 2020; Tasserie et al., 2022; Muller et al., 2023). Importantly, these latter studies suggest distinct roles for different thalamic nuclei, dependent on whether the regions contain many (or few) axons that project widely throughout the striatum and cerebral cortex - i.e., the so-called ‘matrix’ thalamic nuclei. Indeed, a neural mass model much like the one used here - albeit with bespoke matrix-like connections - was able to recapitulate (through simulation) the neural signatures of conscious arousal observed following the stimulation of the central lateral thalamus, an area rich in matrix neurons. For this reason, we anticipate that future studies that incorporate this crucial neuroanatomical feature of the thalamus will provide additional insights into the manner in which the brain supports conscious awareness.

Dexmedetomidine, the anaesthetic used in this study, strongly decreases the noradrenergic tone by acting as a alpha-2A adrenergic receptor agonist that increases auto-inhibition of the locus coeruleus (Sanders & Maze, 2007), a key structure controlling arousal (Wainstein et al., 2022). Consistent with our results, the corticothalamic system is strongly related with the neuromodulatory system, where it has been shown that noradrenergic modulation shapes the function of thalamic cells shifting their firing rate to the tonic modes characteristic of awake states (Rodenkirch et al., 2019). The ascending noradrenergic system is also known to shape arousal, and closely relates to 1/f exponent and alpha oscillations (Pfeffer et al., 2022; Dahl et al., 2022). Moreover, our approximation shows a putative different relation between the 1/f exponent and state of consciousness beyond a modulation of cortical excitatory and inhibitory balance (Gao et al., 2017; Medel et al., 2021; Trakoshis et al., 2020), pointing to thalamocortical gain modulation of aperiodic activity and complex EEG dynamics (Washke et al., 2021; Kosciessa et al., 2021) as a key component for arousal modulation.

Our study, while shedding light on the neural mechanisms behind conscious experience and sensory disconnection during dexmedetomidine sedation, has some limitations. First, our neural mass model can oversimplify the complexities of neurobiology, providing a broad understanding but not capturing the full intricacies of brain interactions. Although this is also a key strength of modelling, whereby the model can be used to determine only those complexities required to drive an observed phenomena. Additionally, our focus on dexmedetomidine sedation may not generalise to other anaesthetic agents, warranting further investigation. Furthermore, our use of scalp EEG data has inherent limitations, including source imprecision and potential artefacts, although muscle artifacts and source solutions are included in the current version of the model (Abeysuriya et al., 2016). Future research should consider employing multimodal neuroimaging with simultaneous scalp EEG to experimentally validate our findings, and provide a more detailed and accurate depiction of neural changes in altered states of consciousness and anaesthesia.

## Methods

### Subjects and Protocol

Twenty subjects were enrolled in the Understanding Consciousness Connectedness and Intraoperative Unresponsiveness Study (NCT03284307). Participants were healthy volunteers between 18 and 40 years old without prior anaesthetic contraindications. Dexmedetomidine was initially administered by a rapid infusion of 3.0 μg kg^-1^ h^-1^ for maintenance. A second step of 10 minutes of infusion of 3.0 μg kg^-1^ h^-1^ followed by a 1.5 μg kg^-1^ h^-1^ maintenance infusion.

Subjects were allowed to rest with their eyes closed for 2-10 minutes. Each rest period was concluded by a researcher calling the participant’s name and initiating a brief structured interview consisting of questions designed to assess if the participant had had conscious experience directly before the name call and if the experience was connected to the environment through the senses (Casey et al., 2022). Two members of the research team evaluated participant answers to code each wake report as connected consciousness (CC; conscious awareness of the environment), disconnected consciousness (DC; a conscious experience but no awareness of the environment, such as a dream), or unconsciousness (Unc; complete lack of experience). Further details of the protocol can be found in Casey et al. (2022).

### EEG acquisition and preprocessing

High-density EEG data were collected using a NA300 EGI system with 256-channel gel caps. Electrodes were manually prepared with the application of electrolyte gel to achieve electrode impedances <50 kΩ. Data were recorded using EGI Net Station Acquisition 5.4 software (Eugene, OR, USA). Preprocessing was done using EEGLAB. Briefly, data were filtered between 0.1 and 55 Hz and visually inspected for noisy channels and epochs, which were removed. Independent component analysis was then computed using the InfoMax algorithm, and components dominated by eye movements or muscle artifacts were rejected. After these cleaning steps, data were averaged and referenced, and the last 20 s of data before the wake report was segmented out for analysis.

### EEG analysis

Power spectra were generated by computing Welch’s method of the Fourier Transform as implemented in NeurDSP (Cole et al., 2019). We parameterized periodic and aperiodic components of the PSD using the FOOOF toolbox (Donoghue et al., 2020), which decomposes the log PSD into a summation of narrowband Gaussian periodic (oscillations) and aperiodic (1/f-like) components for the whole frequency range. We set the algorithm with peak width limits of 1-8, maximum number of peaks: 6; peak threshold 1.5; and ‘fixed’ aperiodic mode. Power spectra were parameterized across the frequency range 1 to 40 Hz. This procedure obtained the alpha relative power (alpha RP) and the 1/f slope of the PSD across the scalp for each state.

### Corticothalamic neural mass

We used the experimentally tested corticothalamic neural mass model as employed in BrainTrack algorithm (Abeysuriya et al., 2005, 2006), which has successfully predicted many features of brain function, including EEG time series and spectra, evoked response potentials, and prevailing arousal state (Rennie et al., 2002; Robinson et al., 2001, 2003; Abeysuriya et al., 2015, 2016). The model describes the aggregate activity of neuronal population in terms of their firing rate ϕ*_a_* and mean membrane potential *V_a_* with *a* ∈ {*e*, *i*, *r*, *s*}. The corticothalamic neural mass encompasses two cortical populations (excitatory *e* and inhibitory *i*) and two thalamic populations (relay *r* and reticular *r*). The membrane potential of a population fluctuates *V* (*t*) as a result of incoming firing rate ϕ_*a*_ (*t*) from other populations and itself according to

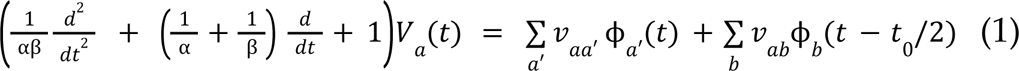

The constants α and β refer to the synaptic decay and rise constants, respectively.*v_aa_* and *v_ab_* correspond to the synaptic strength between populations, where refers to synaptic strength between the thalamic and cortical populations. Axonal propagation between thalamic and cortical populations is delayed by *t*_*a*_. At the cell body, the membrane potential *V*_*a*_ is transformed into a firing rate using a sigmoid function

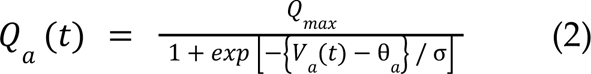

The mean cortical excitatory firing rate is further temporally damped using the following expression:

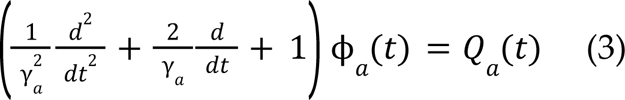

with γ_*a*_ being the temporal damping rate, based on γ = *v*_*a*_ /*r_a_*, where *v*_*a*_ is the propagation velocity and *r*_*a*_ is the mean range of axons.

Equations 1-3 describe the full nonlinear model, which is first transformed to a linear model using linearization around a stable fixed point. Linearization is achieved by expressing the sigmoid function (Equation 2) that transforms *V*_*a*_ (*t*) into *Q_a_* (*t*) as Taylor expansion and retaining only the term containing the first derivative ρ_*a*_ evaluated at the fixed point. Details can be found in Abeysuriya & Robinson, 2016. Using the derivative ρ_*a*_’ the synaptic strengths can be expressed as gain parameters in the linear regime *G_ab_* = ρ_*a*_ *v*_*ab*_. These gain parameters are transformed into three gain parameters describing the cortical (X), corticothalamic (Y), and intrathalamic loop (Z) gains

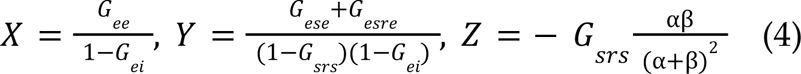

Following this, the linear system in time domain is rewritten in Fourier domain from which it can have an analytical expression for the power spectrum *P*(*w*) as a function of frequency ω

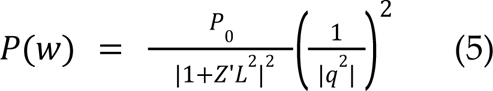

*P*_0_ is a normalisation constant (Robinson et al., 2002) and *q* follows from the dispersion relation described in Abeysuriya & Robinson, 2016,

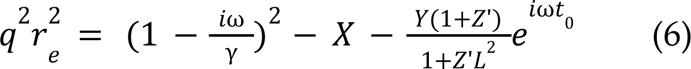

*Z*’ follows from a transformation of Z

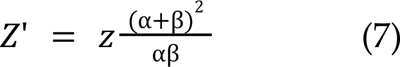

and *L* follows from the transformation of the second-order differential operator describing the synaptic response (Eq. 1) in Fourier domain, which can be interpreted as a low-pass filter depending on the synaptic parameters *α* and *β*

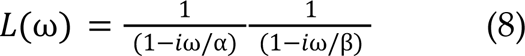

BrainTrack also accounts for pericranial muscles results in electromyogram (EMG) artefacts on the EEG, *P_total_(ω)* = *P(ω)* + *P_EMG_*(ω).

### EEG power spectrum

The model equations (1, 2, …) can be integrated numerically to obtain the EEG time series. By taking the Fourier transform of the simulated EEG signal, we are able to express the firing rate ϕ_*e*_ in terms of the external signal ϕ_*n*_. In particular, we find

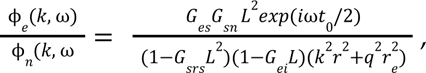

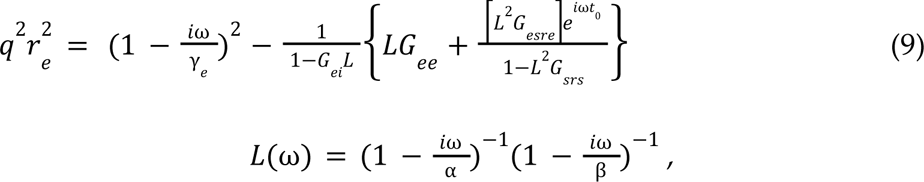

where k and ⍵ are the wave vector (with magnitude k=2π/λ where λ is the wavelength) and angular frequency (⍵=2πf where f is the usual frequency in Hz), respectively. The gain *Gab* = *ρaνab* = *ρaNabsab* is the response in neurons *a* due to unit input from neurons *b*. The quantities *Gese* = *GesGse*, *Gesre* = *GesGsrGre* and *Gsrs* = *GsrGrs* correspond to the overall gains for the excitatory corticothalamic, inhibitory corticothalamic, and intrathalamic loops, respectively. The input stimulus *ϕn* is taken to be white noise

### Brain State tracking

The parameter set x = [*X, Y, Z,* α, β, *t*_0_, *P*_*EMG*_] is is determined through the optimization process of minimizing the discrepancy between the experimentally acquired power spectrum *P_exp_* and the model-generated power spectrum *P_total_* (*x*), expressed as

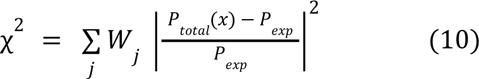

Here, the index j represents the frequency bins, and the weights *W_j_* are designed to provide weighting across frequency decays, being proportional to the inverse of the frequency (1/f). Due to the expansive parameter space, the BrainTrack algorithm constraints parameter values to align with neurophysiologically plausible ranges (Robinson et al., 2004, Abeysuriya et al., 2015, 2016). The χ statistic is subsequently transformed into a likelihood function.

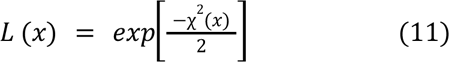

To maximize the likelihood function and minimize error, we employed the Metropolis–Hastings algorithm (Metropolis et al., 1953), which generates probability distributions for each parameter through a Markov chain random walk. Detailed information about this algorithm can be found in Abeysuriya et al., 2016. For each subject, we applied the Metropolis–Hastings algorithm to derive individual power spectra model parameters. The random walk began with initial parameters obtained from a comprehensive database of healthy control subjects (Abeysuriya et al., 2015), primarily influencing the convergence time rather than the final output. At each subsequent step of the random walk, we calculated the likelihood using Eq. 11. A new set of parameters was proposed, and its likelihood was computed similarly. If the new parameters had a higher probability, we accepted the step for sampling the probability distribution. If not, we randomly selected a number from a uniform distribution. If this random number was less than the ratio of the new parameters’ probability to the old parameters, the step was accepted for sampling the probability distribution. Conversely, if the random number was greater than the ratio, the step was not accepted. This iterative process continued until no further changes occurred in the sampled probability distribution.

### Statistics

The relation between 1/f exponent and alpha RP was calculated using nonparametric Spearman rank-order correlation coefficient. Differences between states, for 1/f exponent, alpha RP and neural field model parameters were assessed by using Wilcoxon signed-rank test, which is a non-parametric version of the paired T-test. All reported p-values were corrected by FDR correction.

## Supplementary Figures

**Supplementary Figure 1.**
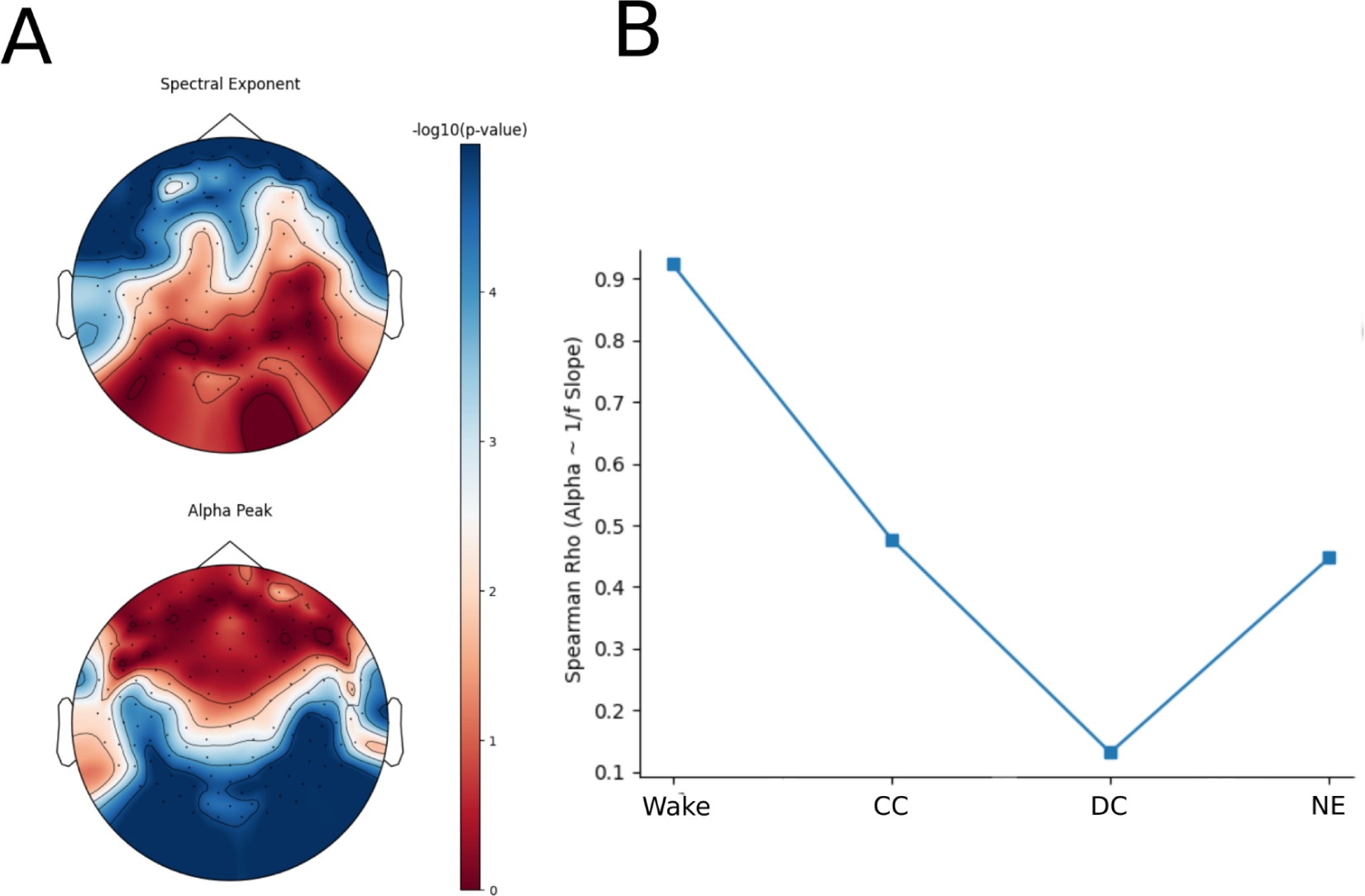
(A) Topographic representation of the correlational structure of alpha relative power and the 1/f spectral exponent. Colorbar represents the -log10 p-value of the Wilcoxon Signed-Rank test between Wake and NE. (B) The spatial correlation (across electrodes) changes depending on the arousal state.

**Supplementary Figure 2.**
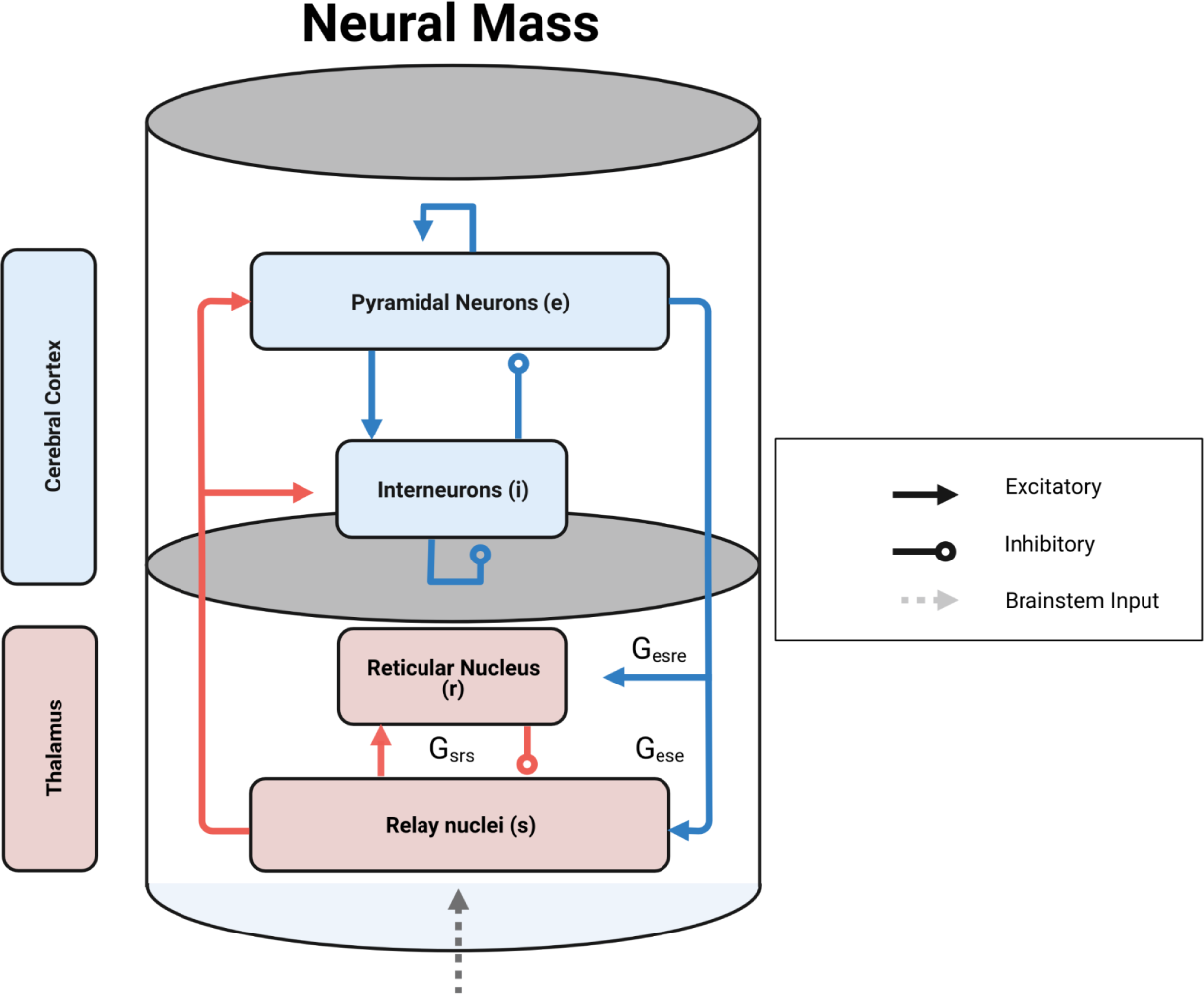
Corticothalamic neural mass model implemented at each node of the network: each mass was comprised of four distinct cellular populations: an excitatory cortical pyramidal cell (“*e*“), an inhibitory cortical interneuron (“*i*”), an excitatory, specific thalamic relay nucleus (“*s*”), and an inhibitory thalamic reticular nucleus (“*r*”), with intranode corticothalamic neural mass coupling defined according to known anatomical connectivity.

**Supplementary Figure 3.**
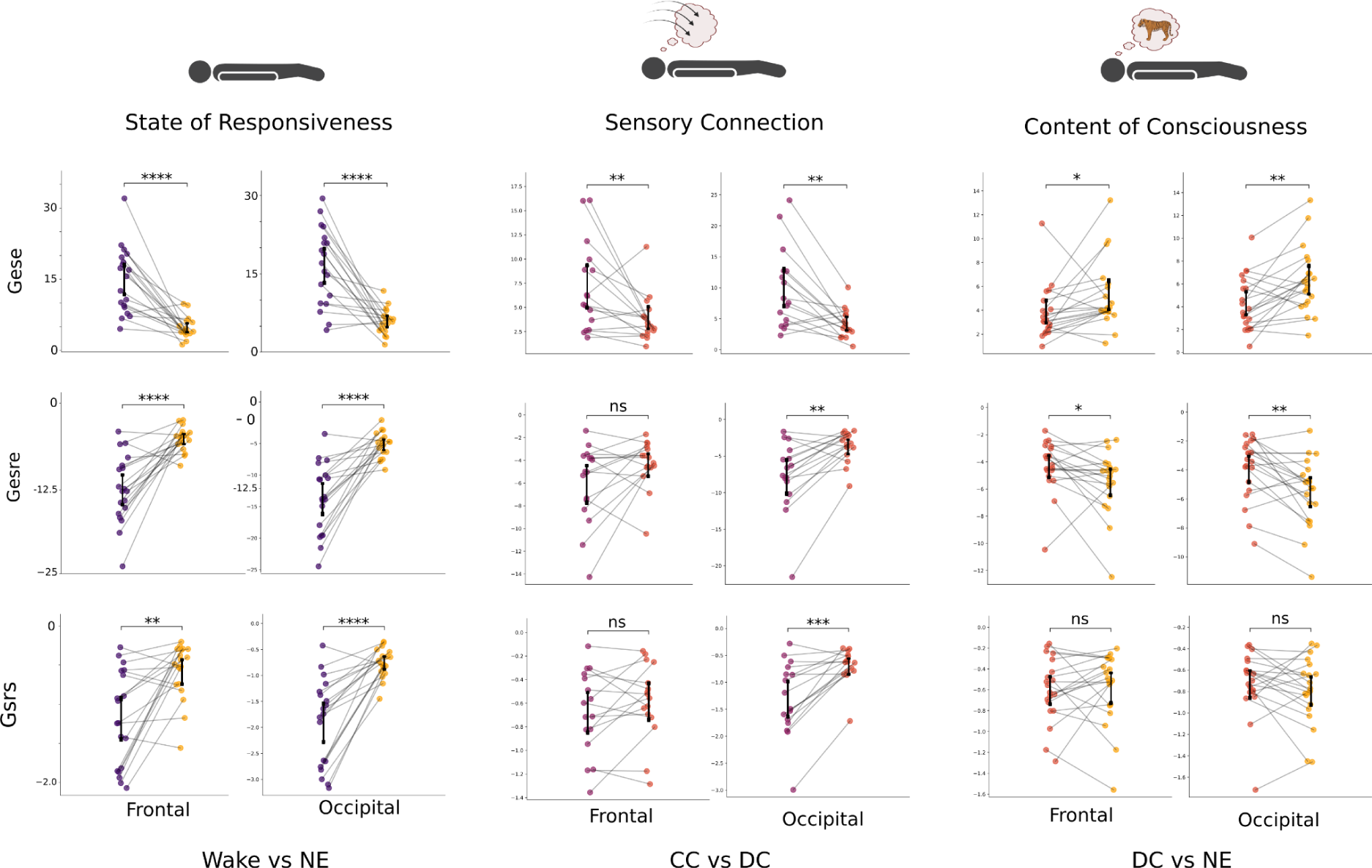
Single comparison for each representative modelling parameter. (* = p<0.05, ** = p<0.01, *** = p<0.005 for FDR-corrected Wilcoxon signed-rank test).

## Notes

### Competing Interest Statement

The authors have declared no competing interest.

