## Supplementary figures and images for "Translating electrophysiological signatures of awareness into thalamocortical mechanisms by inverting systems-level computational models across arousal states"

### Supplementary Figure 1

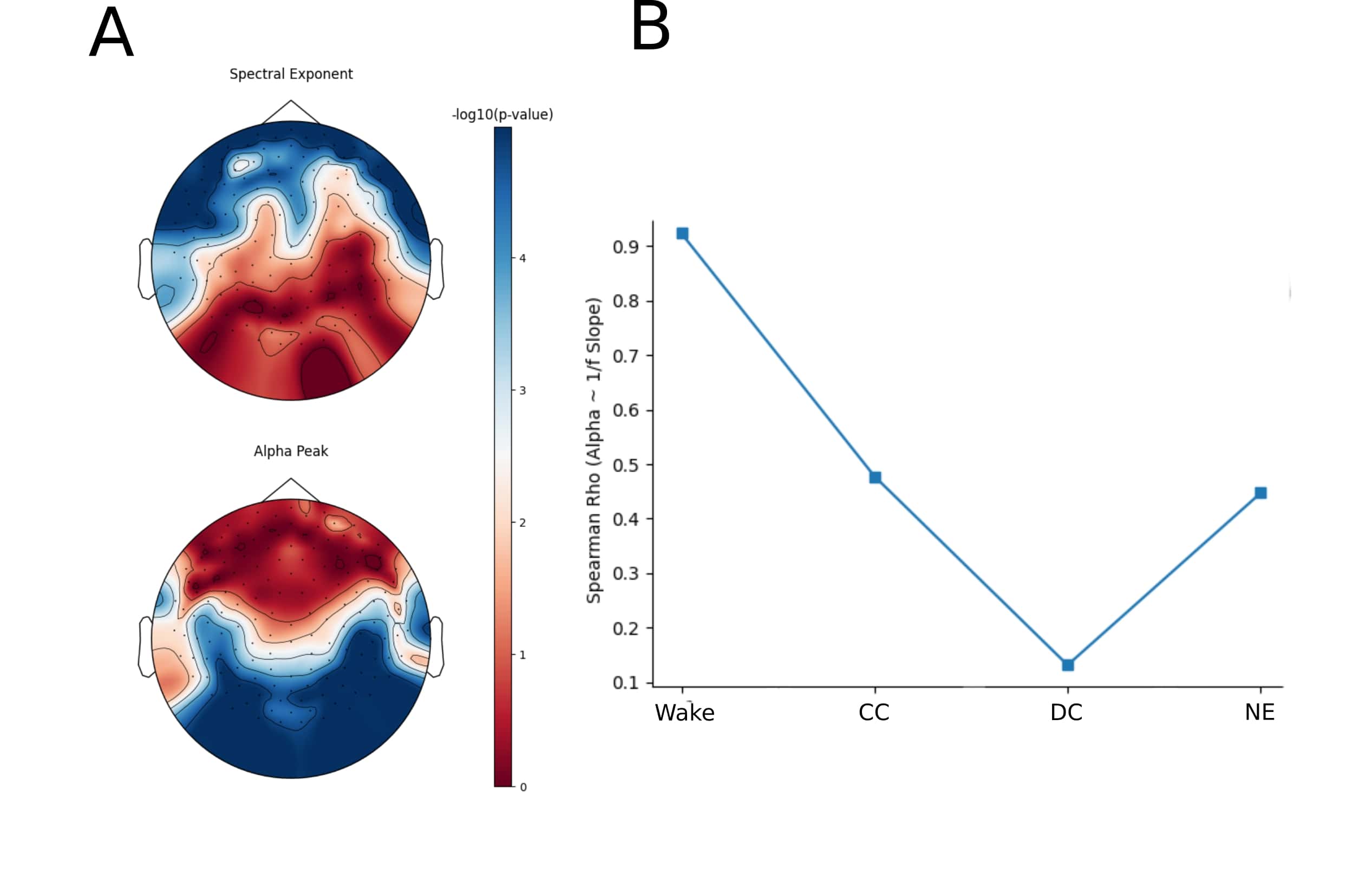

### Supplementary Figure 2

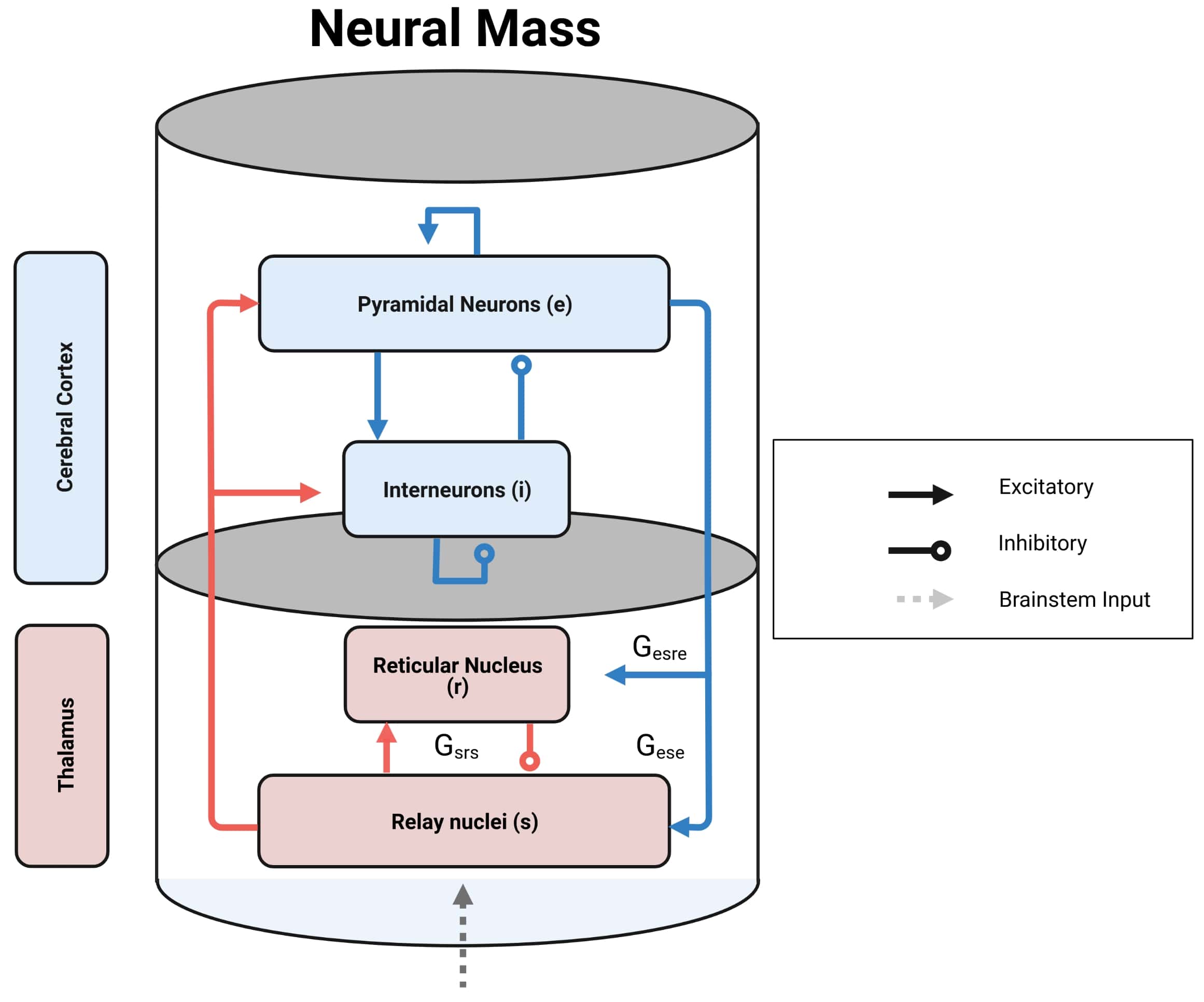

### Supplementary Figure 3

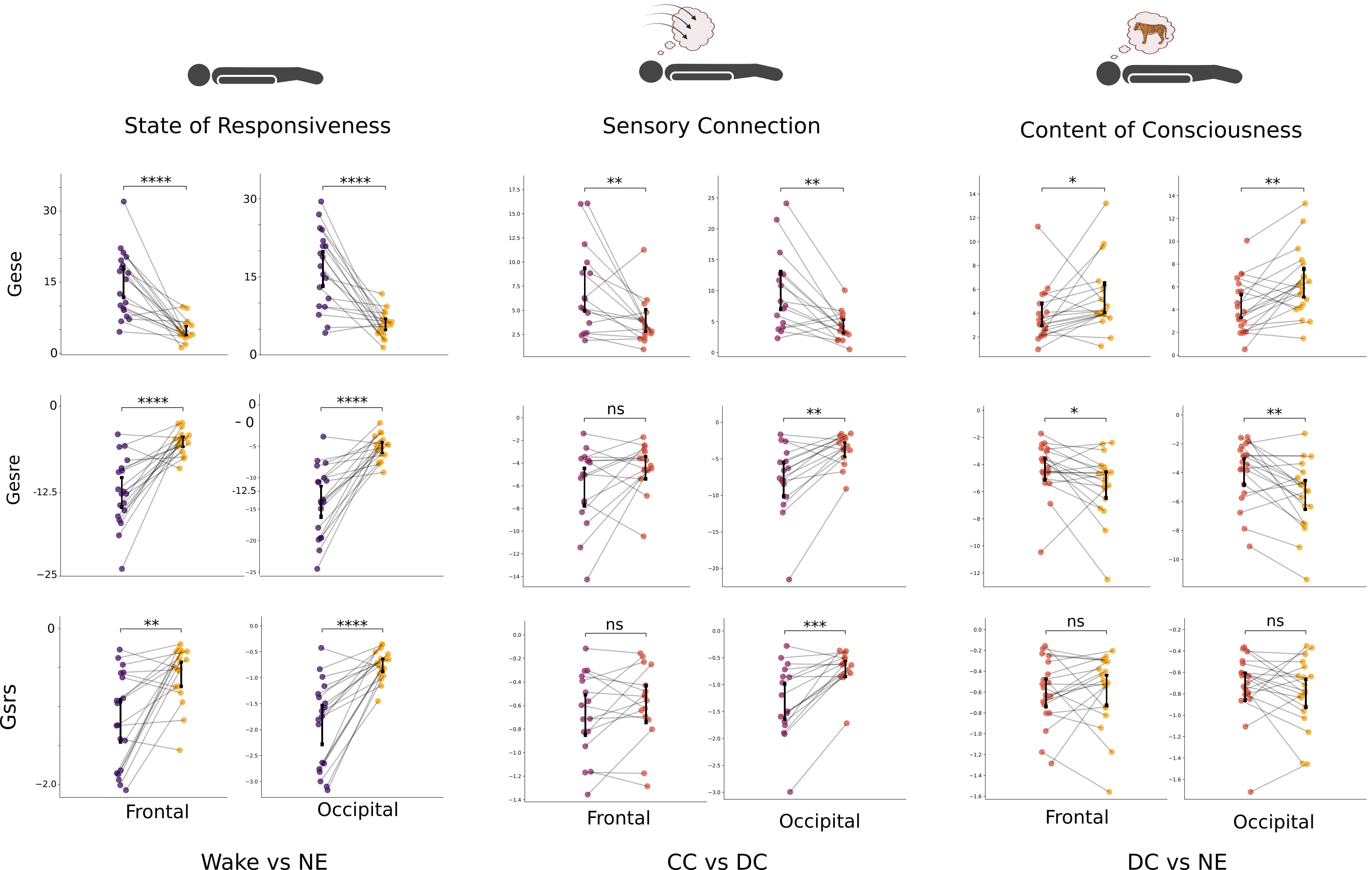
